# MM-MRE: a new technique to quantify individual muscle forces during isometric tasks of the wrist using MR elastography

**DOI:** 10.1101/582825

**Authors:** Andrea Zonnino, Daniel R. Smith, Peyton L. Delgorio, Curtis L. Johnson, Fabrizio Sergi

## Abstract

Non-invasive *in-vivo* measurement of individual muscle force is limited by the infeasibility of placing force sensing elements in series with the musculo-tendon structures. At the same time, estimating muscle forces using EMG measurements is prone to inaccuracies, as EMG is not always measurable for the complete set of muscles acting around the joints of interest. While new methods based on shear wave elastography have been recently proposed to directly characterize muscle mechanics, they can only be used to measure muscle forces in a limited set of superficial muscles. As such, they are not suitable to study the neuromuscular control of movements that require coordinated action of multiple muscles.

In this work, we present multi-muscle magnetic resonance elastography (MM-MRE), a new technique capable of quantifying individual muscle force from the complete set of muscles in the forearm, thus enabling the study of the neuromuscular control of wrist movements. MM-MRE integrates measurements of joint torque provided by an MRI-compatible instrumented handle with muscle-specific measurements of shear wave speed obtained via MRE to quantify individual muscle force using model-based estimator.

A single-subject pilot experiment demonstrates the possibility of obtaining measurements from individual muscles and establishes that MM-MRE has sufficient sensitivity to detect changes in muscle mechanics following the application of isometric joint torque with self-selected intensity.

## I. Introduction

Quantification of individual muscle force applied during tasks that require coordinated muscle co-activation would provide considerable insights in neuromuscular physiology, and enable accurate diagnosis and management of different neuromotor disorders. However, direct measurement of individual muscle force requires to place force sensing elements in series to the musculotendon units. Since such protocol can only be performed using invasive procedures involving muscles dissection [1], currently, direct measurements of muscle force is not possible *in-vivo* in humans.

To solve this problem, over the years, researchers have proposed alternative solutions to quantify individual muscle force, integrating indirect measurement of muscle activity, joint torque and angle into muscle force estimators [2]. There are two traditional estimation approaches: forward dynamics and inverse dynamics. While the inverse dynamic approach has been commonly used to study biomechanics of locomotion [3], it has several disadvantages. Estimators based on inverse dynamics do not use measurements of muscle activity and use optimization to estimate individual muscle force. Since optimization uses cost function whose validity is difficult to assess, inverse-dynamics methods can only provide a rough estimate of the net muscle forces [4].

Forward dynamic estimators, on the contrary, use measurements of muscle activation to solve the redundancy of the musculoskeletal system and, for this reason, they do not require the assumption of any cost function. Typically, the measurement of muscle activation is experimentally obtained using surface electromyography (sEMG) [5], [6]. As sEMG can only measure the activity of superficial muscles, current estimation approaches neglect the contribution from non-superficial muscles, leading to significant error in estimating muscle force [7]. Moreover, since sEMG essentially quantifies the neural drive sent to the muscles, it is insensitive to those factors that define the muscle mechanics such as muscle length, velocity, pennation angle, and fatigue that, then, need to be included in the estimator using models (e.g. Hill type muscle model [8]).

More recently, novel approaches that employ shear wave elastography [9], [10] have been proposed to directly quantify muscle mechanics. The underlying assumption of these techniques is that change in the mechanical load of the musculotendon unit results in a change of the velocity with which shear waves would propagate in the tissue. As such, with proper equipment capable of stimulating and recording wave motion in the tissue, such as shear wave ultrasound elastography machines [9] or custom-developed tendon-tapping devices and accelerometer-based sensors [10], it is theoretically possible to directly quantify muscle load.

While approaches based on shear wave speed measurement have been extensively used in recent studies and have already produced significant advances in our understanding of muscle mechanics during voluntary motor tasks, previous methods can only be used to study a limited set of superficial muscles. As such, these methods are not suitable to study the coordinated motor actions of muscles that are arranged on more layers, such as those in the forearm that control wrist joint motion and fine object manipulation. Among all shear wave elastography techniques, magnetic resonance elastography (MRE) [11]–[13] is the most promising technique to quantify individual muscle force during coordinated wrist and hand movements as it offers excellent properties in terms of tissue penetration and field-of-view size. However, even though MRE has been successfully used to estimate the 3D shear modulus in different tissues of the human body [13], its application to study human skeletal muscles has been limited [14]. Because of temporal limitations (acquisition time ranged from about 30 s for 1D measurements to several minutes for 2D measurements [15]), in fact, current applications of this technique to study contractile states of muscles has been very limited, with few studies that only focused on individual muscles, and not on coordinated muscle function.

In this work, we present multi-muscle magnetic resonance elastography (MM-MRE), a new technique that integrates measurements of the shear wave speed obtained via MRE from all muscles in the forearm with measurements of joint torque and angles measured via an MRI-compatible instrumented handle. A third component of MM-MRE is model-based forward dynamics estimation that integrates measurements from individual muscles with measurements of joint torque, thus allowing estimation of individual muscle force during wrist isometric tasks. The main goal of this paper is to describe methodological aspects of our novel technique mainly focusing on three aspects: the design of a MR-compatible robotic device that allows us to safely measure wrist joint torques at different joint posture during MR imaging; the implementation of an MRE pulse sequence capable of rapidly sampling shear wave displacements throughout all forearm muscles; and the mathematical derivation of the estimator that integrates the measurements of shear modulus and joint torque to return estimates of individual muscle force.

## II. Multi-Muscle Magnetic Resonance Elastography

Applicability of MRE to quantify individual muscle force requires some preliminary considerations about (1) the relationship between the shear wave propagation velocity and the mechanical characteristics (load and stiffness) of the muscle, (2) the relationship between shear and Young’s modulus, and (3) the force-stiffness relationship in skeletal muscles.

First, while for isotropic tissues the shear wave speed squared 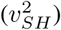 is linearly related to the shear modulus (*G*), in axially isotropic tissues, such as tendon and muscle fibers, shear wave speed depends on both shear modulus and axial load (*σ*). A recent study showed that shear wave propagation in tendon fibers is described by the model of a tensioned Timoshenko beam [10]:

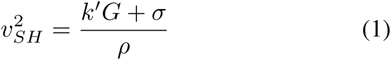

where *ρ* is the density of the muscle and *k′* is the shear correction factor (0 < *k′* < 1).

Second, even though the assumption of a linear relationship between shear (*G*) and Young’s (*E*) modulus is theoretically valid only for isotropic and homogeneous materials, a recent study observed a strong linear relationship between *E* and *G* also for muscle tissue [16], despite being the muscle tissue intrinsically anisotropic:

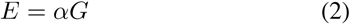

While for isotropic material the proportionality constant is approximated as *α* = 3, Eby *et. al* [16] observed that the value of *α* is muscle-specific and ranges in the interval [4–5].

Lastly, it is well known that skeletal muscles undergoing isometric contractions are characterized by force-dependent stiffness properties. This intrinsic property is called Short Range Stiffness (SRS) [17], [18] and is directly derived from the cross-bridge muscle model, where an increase in the number of actin-myosin cross-bridges results in an increase of muscle axial and shear modulus. A widely accepted model of the SRS assumes that the Young modulus of muscles increases linearly with load:

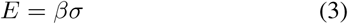

where *β* is a muscle specific proportionality constant.

Given these premises and substituting Eq. 2 and Eq. 3 in Eq. 1, it is possible to observe that there exists a linear relationship between shear wave speed squared and muscle load:

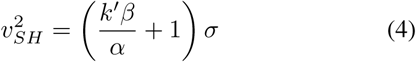

Finally, since the load *σ* is related to the muscle force *f*_*MT*_ through the muscle cross section area (*A*_*CS*_) it is possible to express the relationship between shear wave speed and muscle force as:

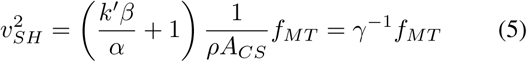

As such individual muscle force can be estimated from measurements of the shear wave speed with a proper calibration procedure capable of determining the set of muscle-specific proportionality constants *γ*_*i*_.

### A. Muscle force estimator

To determine the set of muscle-specific *γ*, we have developed a forward-dynamics muscle force estimator (Fig. 1). The estimator integrates measurements of joint angle, joint torque, and shear wave speed with the forearm geometry obtained from a widely used musculoskeletal model (MSM) [19]. The MSM includes *m*_*tot*_ = 15 muscle spanning the fingers, wrist and elbow joints. For this analysis we considered the upper arm to be grounded, and the hand posture fixed in cylindrical grasp configuration with motion allowed only about the two axes of the wrist joint–i.e. wrist flexion/extension (FE), and wrist radio/ulnar deviation (RUD).

**Fig. 1.**
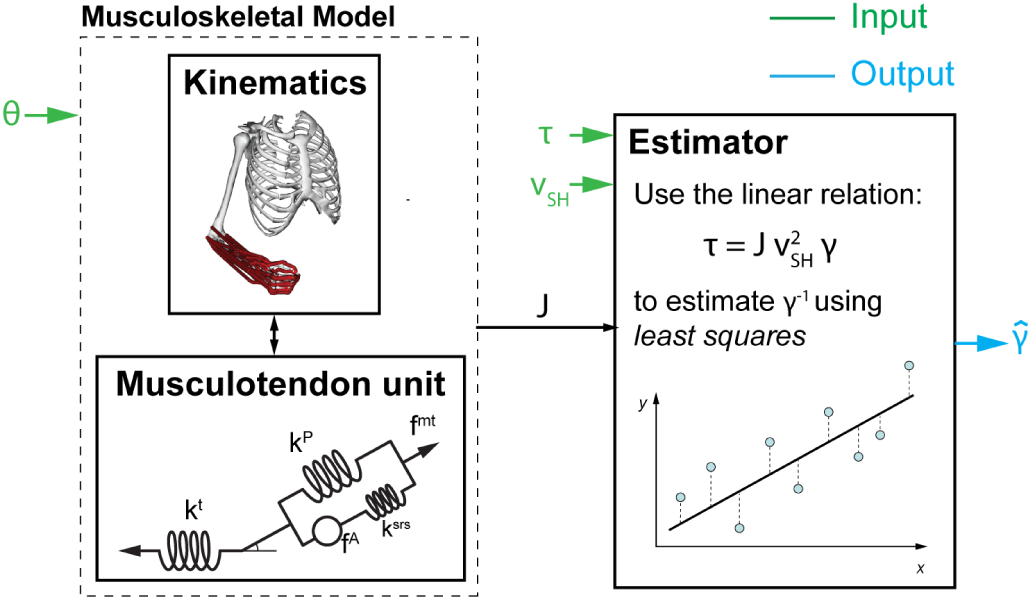
Schematic structure of the proposed estimator.

In this setup the vector of the joint torque measured about the two axes of the wrist joint (***τ*** = [*τ*_*FE*_; *τ*_*RUD*_]) is related to the vector of muscle forces 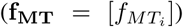 by the equation:

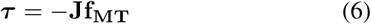

where **J** is the muscular Jacobian whose component *r*_*ij*_ represents the moment arm of the muscle *j* with respect to the joint angle *i*. The bold notation refers to vector and matrices.

Integrating Eq. (5) and Eq. (6), it is then possible to obtain the calibration equation:

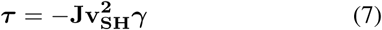

where 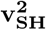 is a diagonal matrix that contains the squared values of shear wave speed measured for each muscle and ***γ*** is a vector that contains the muscle-specific proportionality constants.

Eq. (7) is a linear equation of the form **y** = **X*β***, as such for proper experimental design composed of *n* isometric contractions applied at different postures, it is possible to define an experimental matrix 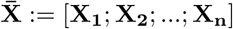 that is of full rank. When this condition is satisfied, it is possible to estimate the vector ***β***:= ***γ*** using a standard least squares fit given measurements of 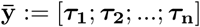.

### B. Model-based experimental design

To verify the existence of a feasible experimental design that allows the estimation of the vector ***γ*** with acceptable error, we have implemented a model-based computational framework that simulates virtual calibration experiments (as described in our previous work [7]). For realistic values of measurement noise and physiological variability 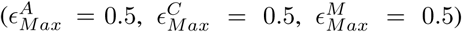 we have simulated six different experimental designs resulting from variation of the joint posture ***θ*** = [*θ*_*FE*_, *θ*_*RUD*_] along one direction (FE or RUD) or two directions (both FE and RUD).

1. ***θ*** ∈ {[−30, 0], [−15, 0], [0, 0], [15, 0], [30, 0]}
2. ***θ*** ∈ {[−30, 0], [0, 0], [30, 0]}
3. ***θ*** ∈ {[0, −30], [0, −15], [0, 0], [0, 15], [0, 30]}
4. ***θ*** ∈ {[0, −30], [0, 0], [0, 30]}
5. ***θ*** ∈ {[0, −30], [−30, 0], [0, 0], [0, 30], [30, 0]}
6. ***θ*** ∈ {[−30, −30], [−15, −15], [0, 0], [15, 15], [30, 30]}

In each design, we assumed that isometric torques are applied in the four cardinal directions (pure wrist FE and wrist RUD torque in both directions), with a magnitude of 1 Nm, followed by a rest condition (zero joint torque). For each experimental design, we then estimated 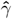 using the proposed estimator, and quantified its accuracy as the percent error in estimating *γ* for the complete set of forearm muscles, averaged across muscles.

As shown in Fig. 2, even though the estimation error is centered at about 10% for all the tested experimental designs, 1D variation of the wrist posture along pure FE is estimated to return the minimal error, while maintaining minimal experiment complexity.

**Fig. 2.**
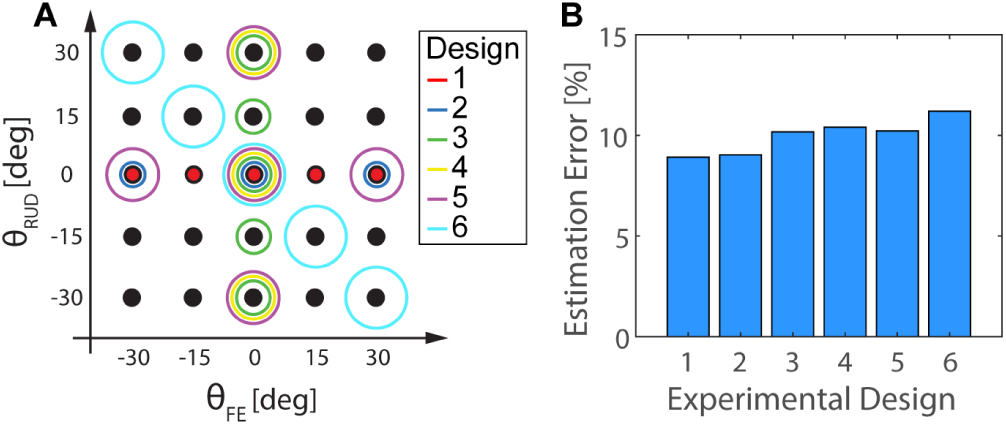
Representation (A) and estimated estimation error (B) for the tested experimental protocols.

### C. Design of the MREbot

We have used the results of the model-based experimental design (Sec. II-B) to inform the design of an instrumented handle, the MREbot, that would be integrated in the MM-MRE methodology. The MREbot features three main components (Fig. 3): a passive locking mechanism that allows the application of isometric wrist joint torques at different wrist postures ([*θ*_*F* *E*_, *θ*_*RUD*_] ∈ {[−30, 0], [−15, 0], [0, 0], [15, 0], [30, 0]}) deg; a support for the distal forearm; a support for proximal forearm that also integrates a support for the MR coil and the drum that applies the vibration required for the MRE protocol (see Sec. II-D).

**Fig. 3.**
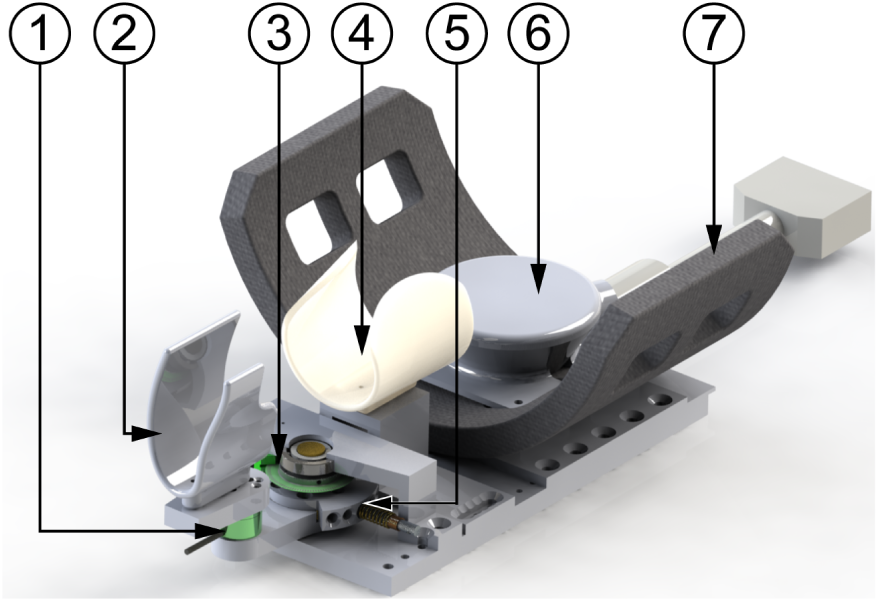
Design of the MREbot. (1) F/T sensor, (2) hand support, (3) optical encoder, (4) forearm support, (5) locking mechanism, (6) mechanical vibrator, (7) MR flex coil.

To ensure MR-compatibility of the entire system, all structural components have been manufactured using ABS-based 3D-printed plastic (RS-F2-GPWH-04, Formlabs Inc., MA, USA) and connected using brass screws, with the output shaft supported by ceramic radial bearings (Boca Bearings, Boynton Beach, FL, USA). Joint posture was monitored using a rotary optical encoder (EM2-1-2500-I, US-Digital, Vancouver, WA), while to measure joint torque we have instrumented the passive mechanism with an MR-compatible F/T sensor (Mini27Ti, ATI Industrial Automation, Apex, NC).

### D. Magnetic Resonance Elastography

MRE is a phase-contrast MRI technique for imaging propagating shear waves [11]. A block diagram describing the different steps of the acquisition and processing of MRE data is shown in Fig. 4. Shear waves are induced by external harmonic vibration of the tissue at a specific frequency. This vibration is synchronized to the MRE pulse sequence that uses oscillating motion encoding gradients to encode the micron-level wave motion into the phase of the MRI signal. Encoding occurs separately in three directions and in time to resolve complex, full vector displacement fields at every point in the imaging volume. In this work we use a single-shot echoplanar imaging (EPI) based MRE sequence to rapidly and robustly capture displacement fields. These displacement fields are then input to a inversion algorithm (NLI) [20] that iteratively solves for the 3D shear wave speed maps.

**Fig. 4.**
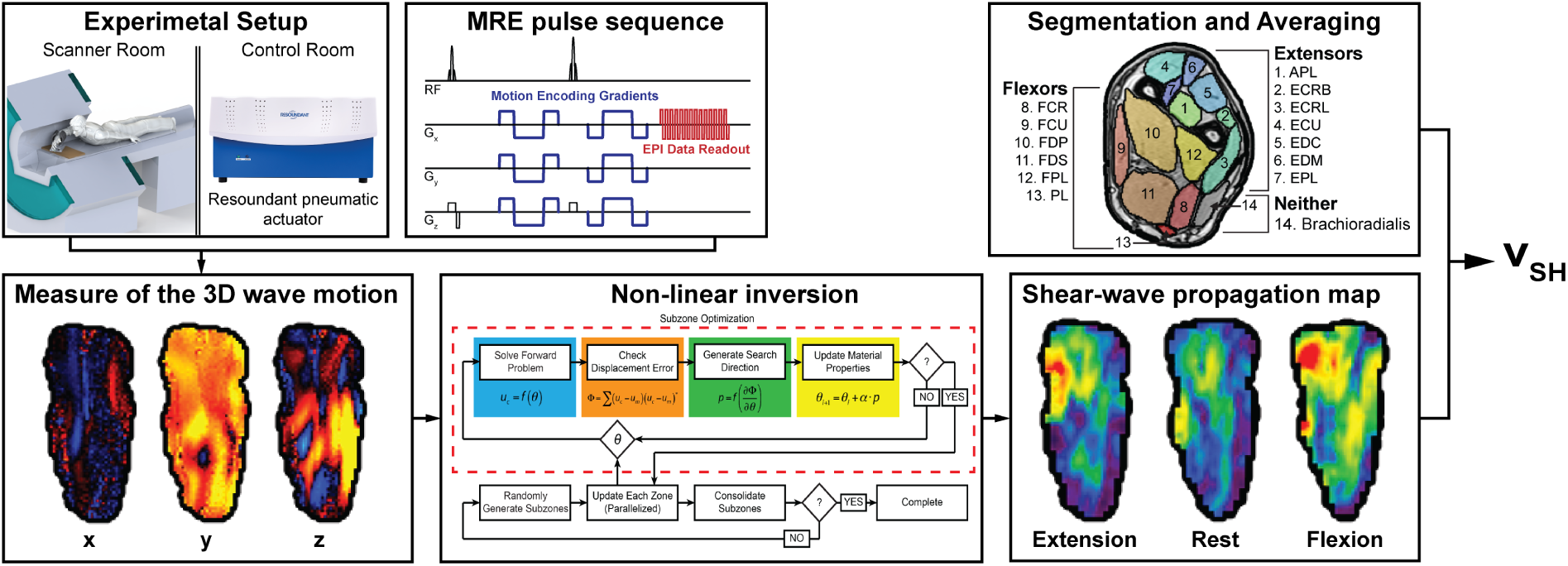
Block diagram of the MRE data acquisition and processing scheme.

Once the 3D maps that quantify the propagation of the shear waves in the forearm have been obtained, muscle-specific measurements of shear wave speed can be obtained by segmenting the maps in different Regions of Interest (ROIs) each of which identifies a different muscle. ROIs are manually identified on T1-weight anatomical scans, which have a greater resolution and contrast than the EPI.

Finally, the outcome measurement of shear wave speed for each individual muscle is obtained by averaging the values of shear wave speed for measured for all voxels in each ROI.

## III. Preliminary Experimental Validation

A preliminary validation experiment was done to demonstrate that during isometric tasks, values of shear wave speed can be extracted for all muscles in the forearm, individually. Moreover, we aimed to establish whether changes in muscle shear wave speed were measurable when self-selected values of flexion and extension torques were applied about the wrist joint. Specifically, we hypothesized that shear wave speed of the flexors would increase during flexion torque, and shear wave speed of the extensor muscles would increase during extension torques.

A single-subject experiment was conducted at the UD Center for Biomedical and Brain Imaging (CBBI) using the Siemens 3T Prisma MRI scanner. In this experiment, one volunteer was instructed to interact with the MREbot and apply isometric contractions. The subject was asked to lay prone with the right arm slightly extended above the head and in the middle of the MRI bore, while wearing the MREbot, and to generate self-selected wrist torques alternating between flexion and extension directions, each lasting 90 s. To minimize muscle fatigue, the experimental protocol was verbally guided so that joint torque was applied only during imaging. A rest period of 30 s was allowed after each contraction. The procedure was repeated 5 times, including a scan in the rest condition to get the baseline levels of shear wave speed after each set of flexion/extension contractions.

MRE scans used an echo-planar imaging (EPI) sequence with 2.0 mm isotropic resolution. Imaging parameters included: TR/TE = 3506/44 ms, FOV = 144 x 240 mm, 72 x 120 matrix, slices = 40. During MRE scans, vibrations were generated at 80 Hz using a pneumatic actuator (Resoundant). An auxiliary isotropic T1-weighted anatomical scan (acquired axially) was included for anatomical localization of individual muscles. The overall scanning time was about 45 minutes. Muscle-specific values of shear wave speed have then been extracted using the methods described in Sec. II-D for the three different contraction states (rest, flexion, extension). Muscle segmentation yieled 13 wrist muscles, plus the Brachioradialis. It is important to note that, while in the forearm, the Brachioradialis does not contribute to either flexion or extension wrist torque.

To test the effects of contraction state on the shear wave speed measured in different muscle groups, we used a 2-way ANOVA with factors “contraction state” (flexion and extension) and “muscle type”. Muscles were classified as flexors or extensors based on the sign of their moment arm in the neutral posture (yielding seven extensors and six flexors). A single outcome measure was extracted for each muscle type by averaging the values of shear wave speed measured from all muscles in the same group.

To test the effects of contraction state on the shear wave speed measured in individual muscles, we used two one-tail paired t-tests to compare the shear wave speed measured during a contraction expected to primarily engage specific muscles (e.g. extension for extensors), with the other two conditions. For the Brachioradialis, two-tail paired t-tests were conducted to quantify the effect of contraction state on *v*_*SH*_. Significance was set to 0.05 type I error rate for all statistical tests.

## IV. Results

Muscle-specific shear wave speed measurements have been extracted from 13 wrist muscles and the Brachioradialis, and are reported in Tab. II, Fig. 5.

**TABLE I.** Anova table

|  | df | SS | MS | F | Prob>F |
| --- | --- | --- | --- | --- | --- |
| Muscle type | 1 | 1.31 | 1.31 | 20.58 | 0.0003 |
| Contraction state | 1 | 0.98 | 0.98 | 15.37 | 0.0012 |
| Interaction | 1 | 1.06 | 1.06 | 16.75 | 0.0008 |

**TABLE II.**
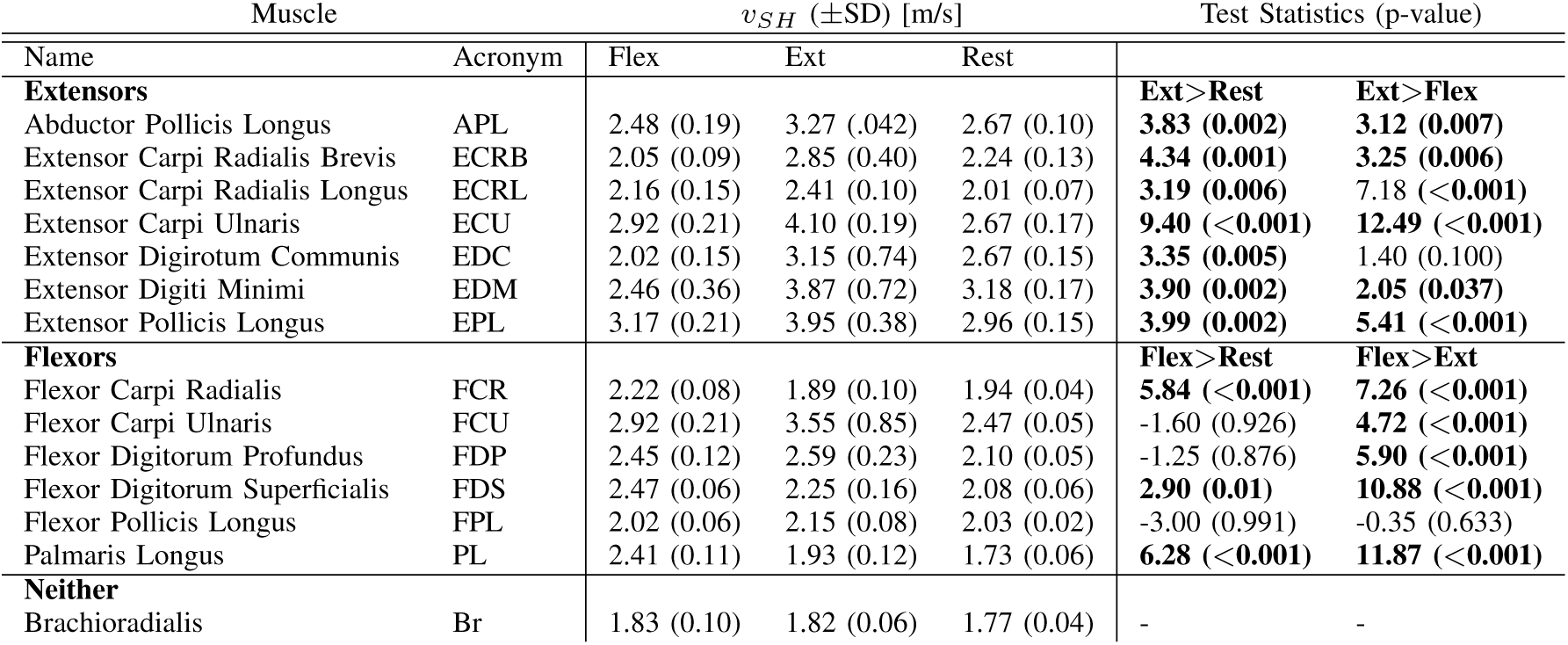
Muscle-specific measurements of the shear wave speed

**Fig. 5.**
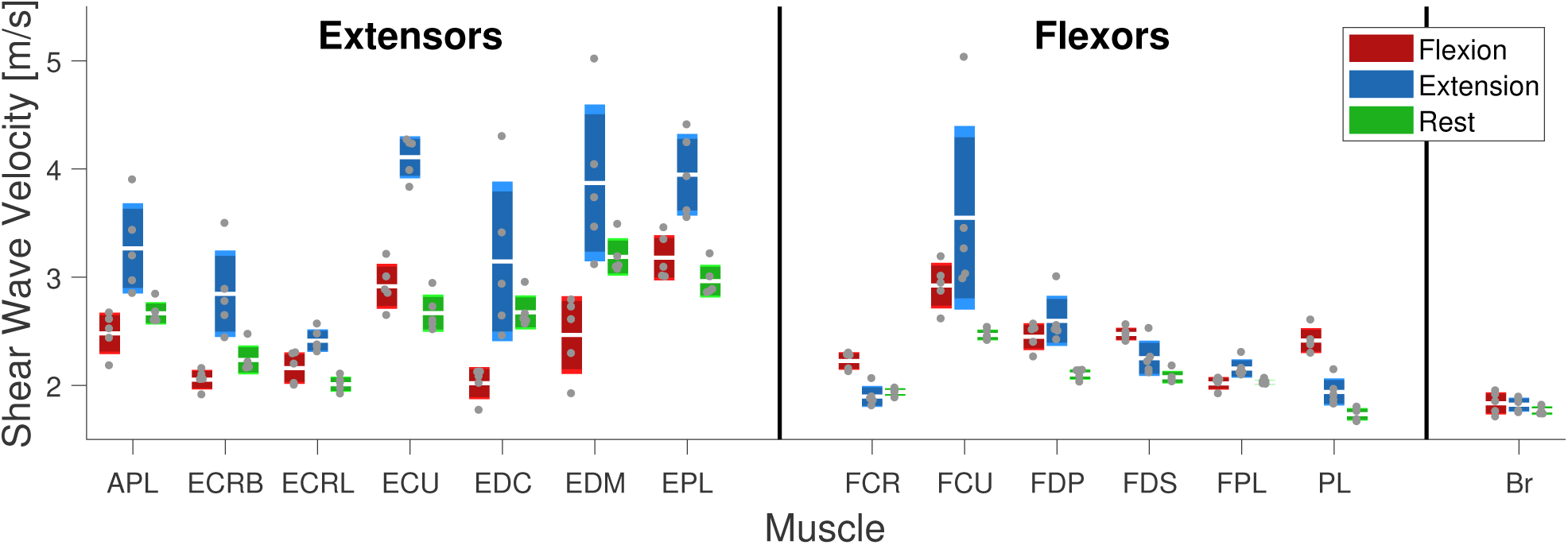
The box plot shows the values of shear wave speed measured for the 14 muscles in the forearm. For each muscle there are three bars that represent the distribution of the measurements obtained in the three different contraction states. In each bar the white line represents the mean of the measurements, the dark box represents the ±standard deviation, with the light box that represents the 95% confidence interval. Finally, the gray dots are the measured values.

Our 2-way ANOVA showed significant effects for both factors–muscle type and contraction state–as well as for their interaction (Tab. III). Post-hoc comparisons show that the shear wave speed was significantly greater in the extensors than in the flexors (mean±SD is 2.92±0.56 m/s for the extensors and 2.41±0.18 m/s for the flexors), and during extension torques than during flexion torque (2.88±0.60 m/s for the extension and 2.44±0.11 m/s for the flexion). Finally, difference in the shear wave speed between flexion and extension was greater for the extensors than for the flexors (0.90±0.46 m/s for the extensors and −0.02±0.27 m/s for the flexors).

Two-sample t-tests (Tab. II) showed that the shear wave speed measured during extension was greater than flexion in all extensors, while it qas greater during extension than during rest in 6/7 extensors. As for the flexors, we can observe a that the shear wave speed is greater during flexion than during extension in 3 out of 6 muscles, while in 5 out of 6 muscles the shear wave speed is greater during flexion than during rest.

For the Brachioradialis, no statistically significant difference can be observed between the three different contraction states.

## V. Discussions and Conclusion

In this paper we have presented and validated Multi-Muscle Magnetic Resonance Elastography (MM-MRE), a novel technique that can be used to quantify forces exerted by all muscles in the forearm during application of isometric wrist torques. MM-MRE combines three key components: i) a fast MRE pulse sequence that allows 3D quantification of the shear wave propagation velocity in a field-of-view that encompasses the entire forearm; ii) an MRI-compatible instrumented handle to record wrist joint torque at different postures during MRE; and iii) a forward dynamics estimator that integrates the measurements of wrist torques and position with muscle MRE data to estimate individual muscle force.

While it was not possible to validate the full protocol due to technical issues with the FT sensor of the MREbot, we conducted a set of experiments to validate key aspects of the MM-MRE protocol. Specifically, our results demonstrate the possibility of obtaining measurements from individual muscles, and establishes that MM-MRE has sufficient sensitivity to detect changes in muscle mechanics that occur as a result of application of isometric torque at the wrist joint.

Our future work will focus on integrating the MREbot within the presented experimental protocol, which will allow to obtain estimates of individual muscle force, and thus investigating the neuromuscular control of coordinated motor action in healthy subjects and neurological patients. Moreover, to achieve the goal of imaging the entire forearm with sufficient spatial resolution during contractions of just 30 s, we plan to separate the x, y, and z encodings with rests in between. This is similar to a “breath hold” protocol employed during MRE scanning of the body portions that move as a consequence of breathing. We will ensure consistent contraction during the three different scans through the FT sensor.

## VI. Acknowledgments

We acknowledge support from the University of Delaware Research Foundation grant no. 16A01402, from ACCEL NIGMS IDeA grant no. U54-GM104941, and from startup funds by the University of Delaware.

